# Transcriptional control of cardiac neural crest cells condensation and outflow tract septation by the Smad1/5/8 inhibitor Dullard

**DOI:** 10.1101/548511

**Authors:** Jean-François Darrigrand, Mariana Valente, Pauline Martinez, Glenda Comai, Maxime Petit, Ryuichi Nishinakamura, Daniel S. Osorio, Vanessa Ribes, Bruno Cadot

## Abstract

The establishment of separated pulmonary and systemic circulations in vertebrates, via the cardiac outflow tract (OFT) septation, stands as a sensitive developmental process accounting for 10% of all congenital anomalies. It relies on the Neural Crest Cells (NCC) colonization of the heart, whose condensation along the endocardial wall forces its scission into two tubes. Here, we show that NCC aggregation starts more pronounced at distal OFT areas than at proximal sites. This spatial organisation correlates with a decreasing distal-proximal gradient of BMP signalling. Dullard, a nuclear phosphatase, sets the BMP gradient amplitude and prevents NCC premature condensation. Dullard is required for maintaining transcriptional programmes providing NCC with mesenchymal traits. It attenuates the adhesive cue *Sema3c* levels and conversely promotes the epithelial-mesenchymal transition driver *Twist1* expression. Altogether, Dullard mediated fine-tuning of BMP signalling ensures the timed and progressive condensation of NCC and rules a zipper-like closure of the OFT.

## Introduction

The heart outflow tract (OFT) is an embryonic structure which ensures the connection between the muscular heart chambers and the embryonic vascular network. Initially forming a solitary tube called truncus arteriosus, it gets progressively remodelled into two tubes which will give rise to the aortic (Ao) and pulmonary (Pa) arteries ((Brickner, Hillis, & Lange, 2000); Figure 1A). This remodelling stands as one of the most sensitive morphogenesis processes. As such, faulty septation of the OFT represents 30% of all congenital heart diseases, with poor clinical prognosis due to improper mixing of oxygenated and deoxygenated blood. This thus calls for a better understanding of the cellular and molecular cues by which the OFT gets septated during development.

**Figure 1:**
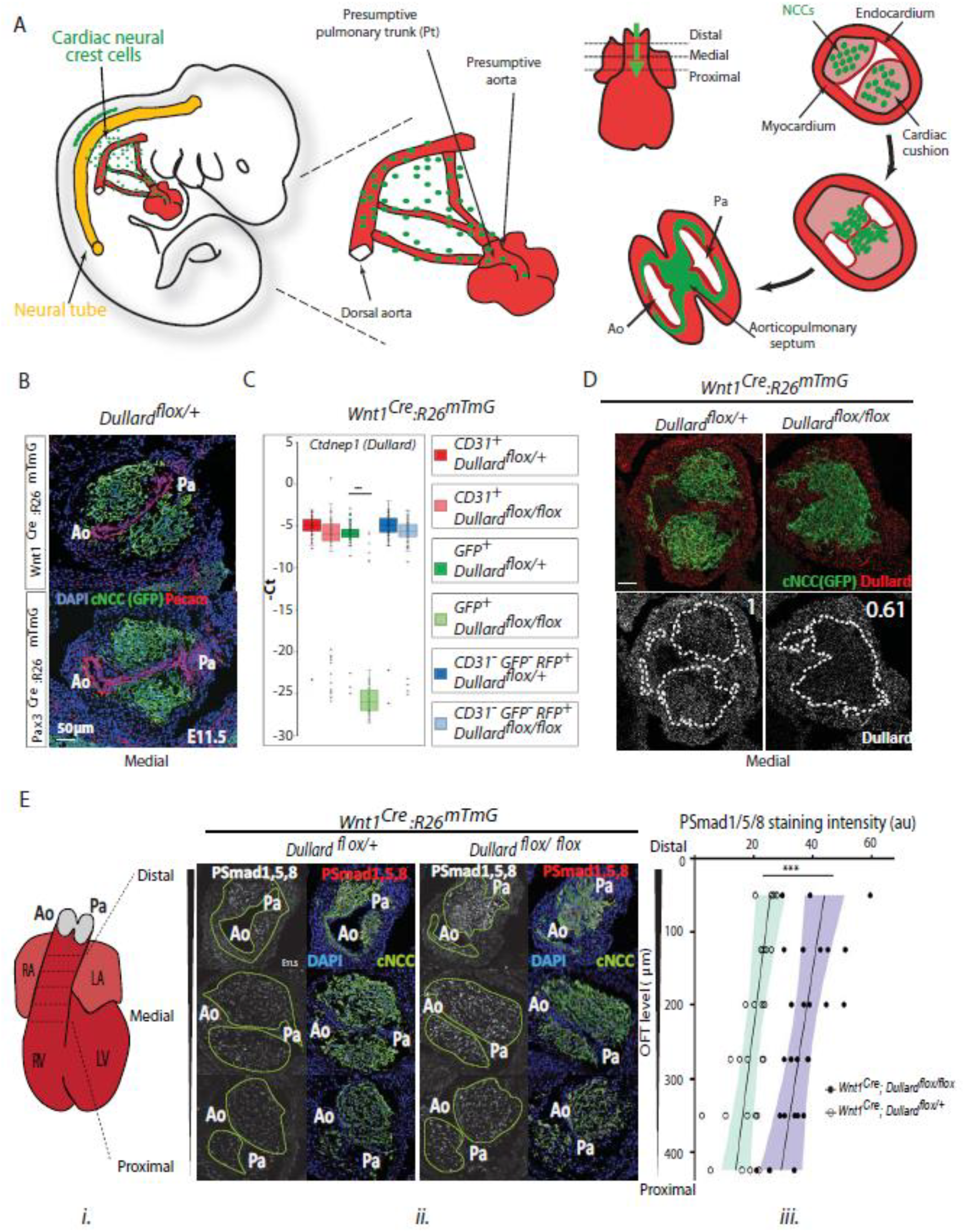
Dullard acts as a Smad1/5/8 activity inhibitor in cardiac NCC. **(A) Ai** Schematic representation of the migration routes the cardiac NCC (green) have taken to reach the heart region (red) in a E10.5 mouse embryo. **Aii** Schematics of the embryonic heart at E11.5 showing the distal-proximal axis of the OFT. **Aiii** Schematic representation of transverse sections through the OFT showing discrete stages of NCC condensation and endocardium septation along the OFT distal-proximal axis. **(B)** Pecam and GFP immuno-labelling and DAPI staining on transverse sections throughout the medial OFT of E11.5 Wnt1^Cre^ or Pax3^Cre^; Dullard^flox/+^; Rosa26mTmG embryos. **(C)** Normalized expression levels of Dullard assayed by q-RT-PCR on single cells isolated from E11.5 Wnt1^Cre^; Dullard^flox/+^ and Wnt1^Cre^; Dullard^flox/flox^; Rosa26mTmG hearts (dots: value for a single cell; boxplot: mean± s.e.m.). The probe monitoring Dullard expression specifically binds to exons 2-3, which are the exons recombined by the Cre recombinase. **(D)** Dullard mRNA distribution detected using RNAscope probes, in transverse sections of E11.5 control and mutant OFTs, assessed by RNAscope. Dullard mRNA levels were significantly reduced in mutant cardiac cushions compared to controls; however, mRNA signals were still detected given the binding of Z pair probes to non-recombined exons 5 to 8 and UTR region. **(E) Ei.** Schematics of E11.5 heart showing the position of the transverse sections used to quantify the levels of the phosphorylated forms of Smad1/5/8 (PSmads) in iii. **Eii.** Immuno-labelling for PSmads and GFP, and DAPI staining on transverse sections across the OFT at 3 distinct distal-proximal levels in E11.5 embryos with the indicated genotype. Pale green dotted lines delineate the area colonized by cardiac NCC. **Eiii.** Quantification of PSmads levels in cardiac NCC along the OFT distal-proximal axis of E11.5 embryos with the indicated genotype (dots: values obtained on a given section; n>4 embryos per genotype recovered from at least 3 liters; the black line is the linear regression, the coloured areas delineate the 95% confidence intervals, ***: p-value < 0,001 for a two way-Anova statistical test). Ao: aortic artery, Pa: pulmonary artery.

The morphogenesis of the OFT is orchestrated in time and space by several cross-interacting cell-types including the myocardial progenitors of the second heart field (SHF), the endocardial cells (EC) delineating the OFT lumen, and the cardiac neural crest cells (cardiac NCC) (Kelly, 2012; Keyte & Hutson, 2012) (Figure 1A). Various genetic manipulations or ablation models have highlighted the predominant role of cardiac NCC in initiating and controlling OFT septation (Bockman, Redmond, Waldo, Davis, & Kirby, 1987; Phillips et al., 2013). Originally, cardiac NCC delaminate from the dorsal neural tube and migrate through the pharyngeal mesoderm to reach the developing OFT (Figure 1A). There, they invade the two cardiac cushions, condense towards the endocardium and trigger its rupture, thereby inducing cardiac cushions fusion and creating the two great arteries (Plein et al., 2015; Waldo, Miyagawa-Tomita, Kumiski, & Kirby, 1998). The rupture of the endocardium is first detected in the regions of the OFT which are the most distal from the heart chambers. In mouse embryos this rupture occurs around 11.5 days of embryonic development (E11.5; Figure 1A) and then expands progressively to more proximal levels. In parallel to these morphogenetic events, NCC will differentiate into the vascular smooth muscles of the aortic arch (Keyte & Hutson, 2012). In addition, NCC will also contribute to the arterial valves (Odelin et al., 2018).

The intense investigations to identify the molecular cues controlling cardiac NCC stereotyped behaviour and differentiation in the OFT mesenchyme have established the importance of the Bone Morphogenic Proteins (BMP), secreted by the outlying myocardium cells from E8.75 onwards (Danesh, Villasenor, Chong, Soukup, & Cleaver, 2009; Jiao et al., 2003; Liu et al., 2004; McCulley, Kang, Martin, & Black, 2008). Indeed, the ablation of Bmpr1a, one of their receptors, or of Smad4 a key downstream transcriptional effectors or conversely the forced expression of Smad7 a BMP signalling antagonist within the NCC lineage all lead to the formation of hypoplastic cushions, a shorter and non-septated OFT, thus phenocopying cardiac NCC ablation experiments (Jia et al., 2007; Stottmann, Choi, Mishina, Meyers, & Klingensmith, 2004; Tang, Snider, Firulli, & Conway, 2010). The knock-out of the ligand BMP4 from the myocardium similarly prevents OFT septation (Liu et al., 2004). However, little is known on the cardiac NCC behaviour and molecular cascades triggered by BMP signalling and responsible for the cardiac NCC mediated OFT septation.

To get insight into these molecular cascades, we decided to dissect the role of Dullard (Ctdnep1) during OFT morphogenesis, a perinuclear phosphatase uncovered as a potential BMP intracellular signalling inhibitor (Sakaguchi et al., 2013; Urrutia, Aleman, & Eivers, 2016). In the canonical BMP signalling cascade, the binding of BMP ligands to their transmembrane receptors leads to the phosphorylation of the transcription factors Smad1/5/8, which translocate to the nucleus and modify the transcriptional landscape of targeted cells (Bruce & Sapkota, 2012). Out of the few cytoplasmic modulators of this phosphorylation identified, including PP1A, PP2B, the inhibitory Smads 6 and 7 and the Ubiquitin degradation pathway, stands Dullard (Bruce & Sapkota, 2012). The Dullard protein is evolutionary conserved from yeast to mammals and expressed in many embryonic tissues, including the developing neural tube and neural crest cells (Sakaguchi et al., 2013; Satow, Kurisaki, Chan, Hamazaki, & Asashima, 2006; Tanaka et al., 2013; Urrutia et al., 2016)(Figure 1D). Several pieces of evidence collected in drosophila, xenopus, and mouse embryos indicate that this enzyme dampens the Smad1/5/8 phosphorylation levels upon BMP stimulation (Sakaguchi et al., 2013; Satow et al., 2006; Urrutia et al., 2016). However, this activity is likely to be tissue specific, as depleting Dullard in the early mouse embryos does not impair BMP signalling, while its depletion in the mouse embryonic kidneys leads to elevated response of cells to BMPs (Sakaguchi et al., 2013; Tanaka et al., 2013). It is worth noting that in these two cases, Dullard appeared as a key regulator of the morphogenetic events regulating the elaboration of embryonic tissues. While it is required early in development for the expansion of extraembryonic tissues, later on it prevents cell death by apoptosis in kidney nephrons (Sakaguchi et al., 2013; Tanaka et al., 2013).

We showed here that deletion of *Dullard* in the cardiac NCC increases Smad1/5/8 activity, leading to premature and asymmetric septation of the OFT and pulmonary artery closure. This BMP overactivation in the cardiac NCC triggers the downregulation of mesenchymal markers (*Snai2*, *Twist1, Rac1, Mmp14* and *Cdh2*) and the upregulation of *Sema3c*, associated with premature cardiac NCC condensation to the endocardium. Our data converge to a model whereby graded BMP activity, *Sema3c* expression and cardiac NCC condensation along the OFT axis set the tempo of its septation from its distal to its proximal regions. Hence, our findings reveal that fine tuning of BMP signalling levels in cardiac NCC orchestrates OFT septation in time and space.

## Results

### *Dullard* deletion triggers hyper-activation of BMP intracellular signalling in cardiac NCC

In order to ablate Dullard in cardiac NCC, we crossed mice carrying floxed alleles of *Dullard* with mice expressing the Cre recombinase from the *Pax3* or *Wnt1* loci (Danielian, Muccino, Rowitch, Michael, & McMahon, 1998; Engleka et al., 2005; Sakaguchi et al., 2013). Cell lineage tracing was achieved by using a ubiquitous double-fluorescent Cre reporter allele, *Rosa26mTmG*, in which Cre-mediated recombination labels the cells with membrane-targeted GFP (Muzumdar, Tasic, Miyamichi, Li, & Luo, 2007). The pattern of cell recombination in the cardiac cushions of E11.5 control embryos carrying either Cre drivers matched with the pattern of colonizing cardiac NCC described by previous lineage analyses (Figure 1B) (Brown et al., 2001; Jiang, Rowitch, Soriano, McMahon, & Sucov, 2000). RT-qPCR on single Fac-sorted (FACS) cells from dissected OFT and RNAscope *in situ* hybridization on histological sections were used to monitor *Dullard* expression (Figure 1C,D). At E11.5, *Dullard* was ubiquitously expressed in all OFT layers of control embryos. In recombined *Wnt1^Cre^; Dullard^flox/flox^; Rosa26mTmG*, the cardiac NCC displayed a strong reduction in *Dullard* levels, while the surrounding tissues remained *Dullard* positive. Strikingly, in these mutants, the NCC formed a unique mass at the distal part of the OFT, while two distinct NCC cushions were present in the control embryos (Figure 1D,E, 2A,B), indicating that Dullard regulates the spatial organization of NCC in the OFT (see below).

We next assessed the relationship between Dullard and the activity of the intracellular effector of BMP signalling, i.e. the Smad1/5/8 (Smads) transcription factors, in mammalian cells. As previously shown (Satow et al., 2006) in the myogenic cell line C2C12, overexpression of Dullard strongly decreased the levels of phosphorylated Smads (PSmads) induced by BMP2 treatment (Figure supplement 1A). In addition, by generating a version of Dullard carrying a phosphatase dead domain, we could show that this inhibitory role relied on Dullard phosphatase activity (Figure supplement 1A). In cardiac NCC, the deletion of *Dullard* was sufficient to double the levels of PSmads within these cells, whatever their position along the distal-proximal OFT axis of E11.5 hearts (Figure 1E - Figure supplement 1B). Hence in this embryonic cell lineage, Dullard also acts as a BMP intracellular signalling inhibitor. Interestingly, in both control and mutant contexts, PSmads levels were more elevated distally than proximally (Figure 1Eiii - Figure supplement 1B), indicating that cardiac NCC harbour a graded BMP response, which declines as they colonize more proximal OFT areas. Dullard in cardiac NCC is required to dampen the magnitude of the BMP signalling response gradient along the OFT length, but does not control its establishment (Figure 1Eiii).

### Dullard is a key modulator of cardiac NCC mediated OFT septation

We next sought to examine the morphology of the OFT in control and *Dullard* mutants, knowing that cardiac NCC controls its septation (Bockman et al., 1987; Phillips et al., 2013; Plein et al., 2015). For this, we monitored, using 3D lightsheet and confocal microscopy, E11.5-12 hearts labelled for the arterial marker, Pecam (Figure 2A,B - Figure supplement 2E, Movie supplement 1, 2). At distal levels, the OFT of control embryos displayed symmetrical septation with two great arteries of similar size (Figure 2Ai). In contrast, *Pax3^Cre^ or Wnt1^Cre^; Dullard^flox/flox^* embryos exhibited asymmetric breakdown of the endocardium on the pulmonary side with obstruction of the pulmonary artery (Pa) (Figure 2Aiv - Figure supplement 2E). At more medial levels in control embryos, the aortic and pulmonary endocardium regions were connected and attached to the presumptive pulmonary valve intercalated-cushion (PV-IC) (Figure 2Aii,iii). Conversely in mutants, this attachment was absent, and the endocardium was displaced towards the aortic side, highlighting premature cushions fusion and OFT septation (Figure 2Av, vi - Figure supplement 2E).

**Figure 2:**
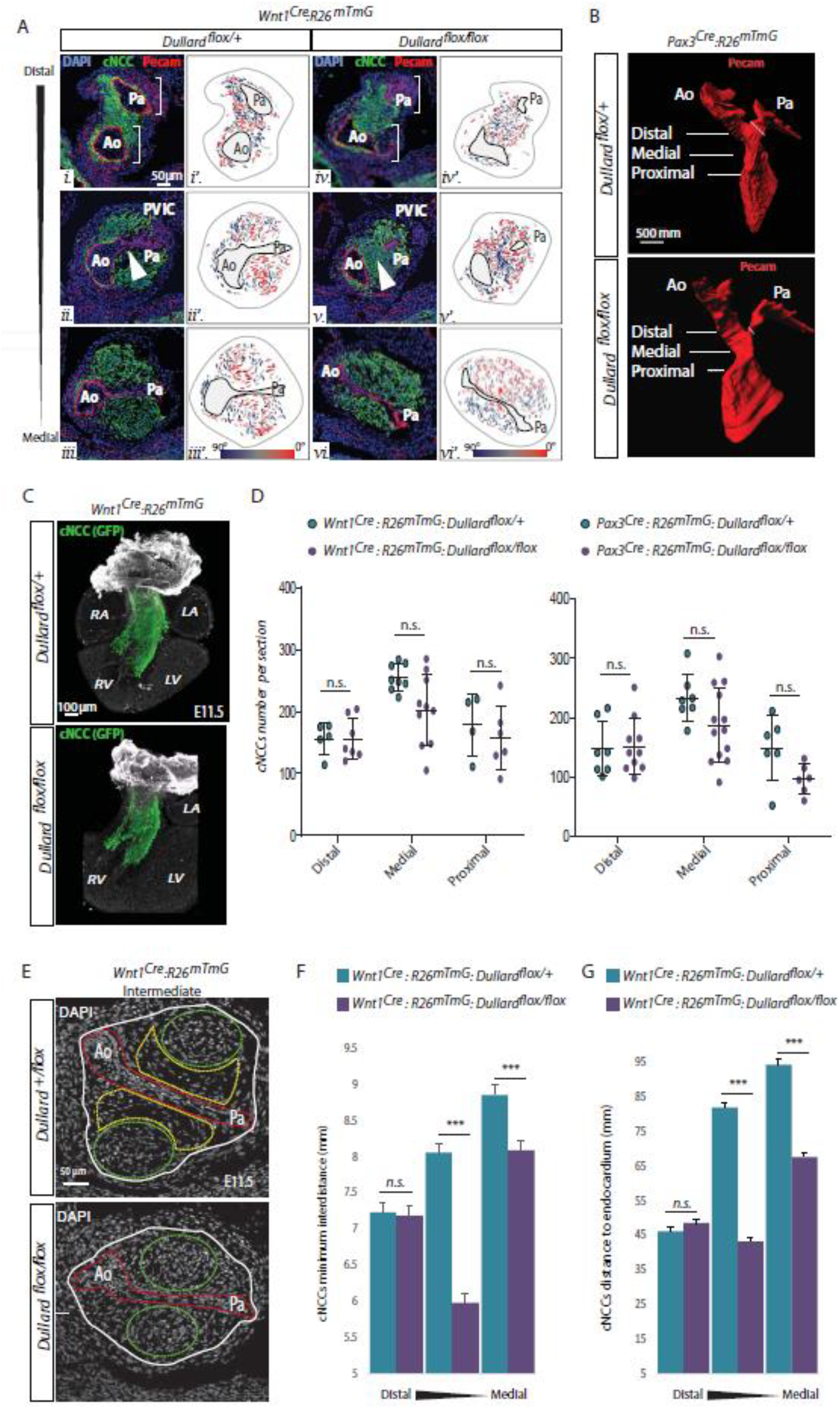
Dullard deletion in cardiac NCC causes asymmetric and premature OFT septation. **(A) Ai-vi.** Immuno-labelling for Pecam (red) and GFP (green) and DAPI (bleu) staining on transverse sections along the distal-proximal axis of the OFT in E11.5 embryos with the indicated genotypes (n>10 embryos collected from more than 3 liters). Brackets in i and iv highlight the symmetric and asymmetric Ao and Pa poles in control and mutant embryos, respectively. Arrowheads in ii and v point at the unruptured and ruptured endocardium in control and mutant embryos, respectively. **Ai’-vi’.** Orientation of the major axis of NCC cells relative to Ao-Pa axis colour-coded as indicated in the section shown in Ai-vi. **(B)** Three-dimensional rendering of the Pecam^+^ endocardium of E12 Pax3^Cre^; Dullard^flox/+^ and Pax3^Cre^; Dullard^flox/flox^ embryos after 3Disco clearing and Lightsheet acquisition (n=3 per genotype). The fine oblique white line marks the Pa width. The OFT levels along its distal-proximal axis analyzed in A are also indicated. **(C)** Three-dimensional rendering of cardiac NCC (green) over Pecam (white) after BABB clearing and confocal acquisition of whole E11.5 hearts isolated from embryos with the indicated genotype (n=2 per genotype). **(D)** Quantification of the total number of GFP^+^ cardiac NCC per OFT section at the indicated distal-proximal OFT axis levels (dots: value per section; bars: mean±s.d.; n.s.: non-significant differences evaluated using a Student t-test). **(E)** DAPI staining on transverse sections through the medial part of the OFT of E11.5 embryos with the indicated genotype. The entire OFT is circled with a while line. The endocardium is delineated in red, the condensed and round NCC in green, the loose and elongated NCC in yellow. **(F)** Minimum distances between NCCs and **(G)** distances between NCCs and the Ao-Pa axis quantified along the distal-proximal axis of the OFT in E11.5 embryos with the indicated genotypes (n=3 embryos from distinct liters were analyzed for each genotype and OFT level, bars: mean±s.d.; ***: p-value < 0.0001 for Student statistical t-test). Ao: Aorta; Pa: pulmonary artery; LA: left atrium; LV: left ventricle; PV-IC: Pulmonary valve intercalated-cushion. RA: right atrium, RV: right ventricle.

To dig into the cellular mechanisms by which Dullard prevents the appearance of these heart defects, we evaluated the migrative, proliferation and death status of cardiac NCC. 3D imaging of these cells thanks to the GFP reporter revealed that cardiac NCC reached similar OFT levels in E11.5 control and *Dullard* mutants, showing that Dullard is not required for NCC colonization of the OFT (Figure 2C - Figure supplement 2A, Movie supplement 2-6). Similarly, staining and quantifying cells in mitosis or in apoptosis using antibodies raised against the phosphorylated form of histone H3 and the cleaved version of Caspase 3, respectively, indicated that Dullard does not control the amplification nor the survival of cardiac NCC (Figure supplement 2B,C). In agreement with these observations, the total number of GFP^+^ cells colonizing the OFT in mutant embryos was not significantly different from that found in controls (Figure 2D).

Finally, we wondered whether the morphogenetic defects of the mutant OFT could stem from differences in cell-cell arrangements, looking at the position and orientation of DAPI labelled NCC and endocardium nuclei (Figure 2Ai’-vi’,E-G). Quantification of the shortest distance between cardiac NCC nuclei indicated that in E11.5 control hearts, NCC condensation was variable along the distal-proximal axis of the OFT (Figure 2E,F). Cells were closer to each other at distal levels than in proximal regions. This progression of NCC condensation along the OFT axis was impaired in *Dullard* mutants. Mutant NCC prematurely condensed notably within the medial region of the OFT (Figure 2E,F). Similarly, the position of NCC to the endocardium was variable along the OFT axis of control embryos; NCC were closer to this epithelium at distal levels than at proximal levels (Figure 2E, G). In mutants, the NCC were in a closer vicinity of the endocardium than control cells, so that in medial levels they displayed traits of cells normally found at distal levels in control hearts (Figure 2E,G). In an agreement with these data, the OFT area was reduced in mutants and remained constant along the distal to proximal axis (Figure supplement 2D). Finally, the orientation of the cardiac NCC nuclei relative to the endocardium appeared also spatially regulated along the proximal-distal axis of the OFT, in both mutant and control hearts (Figure 2Ai’-vi’,E - Figure supplement 2E). In controls, at distal levels NCC were perpendicular to the endocardium, while at proximal levels no orientation preference could be assigned (Figure 2Ai’-ii’, E, Figure supplement 2E). Strikingly, in *Wnt1^Cre^; Dullard^flox/flox^* OFTs the perpendicular orientation was more widely observed than in control OFT (Figure 2Aiv’-vi’,E - Figure supplement 2E).

Altogether these data demonstrate that *Dullard* stands as a key modulator of NCC behavior dynamics in the heart and hence of OFT septation. It hampers NCC condensation, and thereby leads to the premature breakage of the endocardium and to obstruction of the pulmonary artery.

### *Dullard* cell-autonomously controls cardiac NCC mesenchymal transcriptional state

To decipher the molecular basis of the defective OFT remodeling observed in mutants, we micro-dissected five E11.5 control and Dullard mutant heart OFTs and sorted the cardiac NCC (GFP^+^) and endocardial cells (CD31^+^;RFP^+^) from the other OFT cell-types (CD31^−^;RFP^+^) (Figure supplement 2F). We then performed single cell RT-qPCR for 44 genes implicated in epithelial-mesenchymal transition (EMT), migration and/or specification of the different OFT progenitor subtypes (Table S1).

T-statistic Stochastic Neighbor Embedding (t-SNE) was first used to plot the distances existing between the 44 gene-based-transcriptomes of individual cells (Figure supplement 3B). It revealed that the 44 chosen genes were sufficient to segregate the 3 isolated cell subtypes (GFP^+^ NCC, the RFP^+^;CD31^+^ endocardial cells and the rest of RFP^+^;CD31^−^ OFT cells)(Figure supplement 3B), validating our approach. Unsupervised hierarchical clustering analysis of all cells refined this segregation and identified six distinct groups of OFT cells (Figure supplement 3C). Importantly, some of these groups contained both control and *Dullard* mutant cells (Groups 1, 3, 5) meaning that their 44 gene-based-transcriptome was not regulated by Dullard. Instead, other groups were enriched for cells with a given genotype (Groups 2, 4, 6), hence harboured a Dullard dependent transcriptional state. Importantly, the groups 1, 3, 5 barely contained GFP^+^ NCC. For instance, the groups 1 and 5 were composed of RFP^+^;CD31^+^ (*Flt1*^+^;*Kdr*^+^*;Nfatc*^+^;*Tek*^+^) endocardial cells and RFP^+^;CD31^−^ (*Tcf21*^+^*;Wt1*^+^) epicardial cells, respectively. In contrast, the groups 2, 4, 6 were enriched for GFP^+^ NCC. This suggests that Dullard acts cell autonomously, its deletion alters the transcriptional states of NCC and barely modulates that of other cell types present in the developing E11.5 heart.

We further characterised the transcriptomic changes underpinning the segregation between mutant and control cardiac NCC. Unsupervised hierarchical clustering analysis of the GFP^+^ NCC transcriptomes showed that the transcriptional state of these cells was variable both in control and in mutant NCC. Five sub-populations of cells could be identified based on their gene expression signature (Sub Pop1 to 5) (Figure 3A-C - Figure supplement 3D), each of them containing an unbalanced ratio of mutant versus control cells (Figure supplement 3F), suggesting that Dullard influences the fate of all NCC subtypes. 2-dimensional representation and visualization of cells on diffusion maps confirmed that the variation in gene expression in NCC was primary associated with the loss of Dullard (Figure 3B,C - Figure supplement 3D-G).

**Figure 3:**
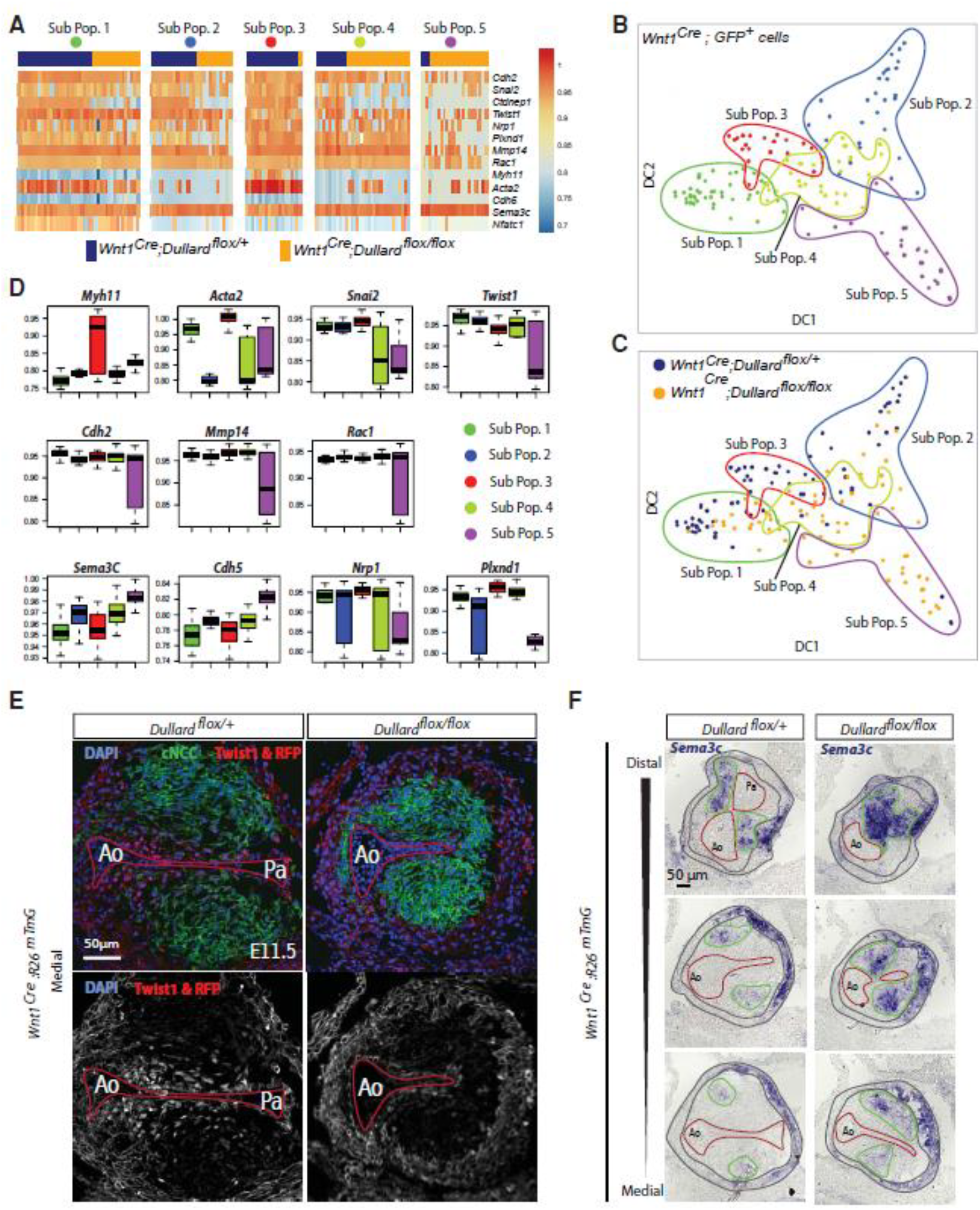
Dullard prevents NCC to acquire a epithelial-like transcriptional states and prolongs the expression of mesenchymal drivers. **(A)** Heatmaps showing the levels of expression of selected genes in all GFP+ NCC in the 5 sub-populations identified using unsupervised hierarchical clustering (cf. Figure supplement 3D) coming from control (blue) or mutant (orange) E11.5 embryos. **(B, C)** Projected position of 44 genes-based transcriptomes assessed in mutant and control cardiac GFP+ NCC on diffusion maps made using the first two Diffusion Component 1 (DC1) and 2 (DC2). In B the 5 sub-populations defined in Figure supplement 3D or in C the mutant and the control cells are highlighted. **(D)** Boxplot representation of the expression levels of genes differentially expressed between the 5 NCC sub-populations (Sub-Pop1: 44 cells, Sub-Pop: 29 cells, Sub-Pop3: 20 cells, Sub-Pop 4: 34 cells, Sub-Pop 5: 24 cells)(mean± s.d.) **(E)** Twist1 immunolabelling and RFP signal (red; grey), as well as GFP (green) immuno-labelling and DAPI staining on transverse sections through the medial OFT of E11.5 Wnt1^Cre^; Dullard^flox/+^; Rosa26mTmG and Wnt1^Cre^; Dullard^flox/flox^; Rosa26mTmG embryos. The red lines mark the endocardium. **(F)** Sema3c expression assessed by ISH on transverse sections from distal to medial OFT levels of E11.5 Wnt1^Cre^; Dullard^flox/+^ and Wnt1^Cre^; Dullard^flox/flox^ embryos. The endocardium is delineated with a red line, the cardiac NCC areas with a green line and the myocardium with grey lines. Ao: Aorta; Pa: pulmonary artery

The NCC Sub Pop1 was characterized by the expression of *Nfatc1, Tcf21, Postn (Periostin)* (Figure supplement 3D,G), which are all expressed in the heart valves (Acharya, Baek, Banfi, Eskiocak, & Tallquist, 2011; Norris et al., 2008; Wu et al., 2011), a heart structure that is also colonized by NCC (Odelin et al., 2018). The slight enrichment for control cells in this sub population rise the possibility that Dullard is required to favour the contribution of NCC to this lineage (Figure supplement 3F). The second population was defined by the predominant expression of cardiac progenitor markers *Tbx1*, *Six2, Gja1, Isl1*, and contained both control and mutant cardiac NCC (Figure supplement 3D,G), indicating that Dullard is not required for the entry of NCC into the muscle lineage (Zhou et al., 2017). Conversely, the emergence of the SubPop 3 which contained NCC more differentiated into smooth muscle cells expressing *Myh11* and *Acta2* (Huang et al., 2008) relies on the activity of this phosphatase; it barely contained *Dullard* mutant cells (Figure 3A-D - Figure supplement 3F). The mutant cardiac NCC clustered in Sub Pop 4 and even more strongly in Sub Pop 5 (28 % of mutant versus 4% of control cardiac NCC) (Figure 3A - Figure supplement 3F). Generally, the genes which were differentially expressed between the Sub Pop 5 and 3, were also altered in Sub Pop4. Yet, the variations were less severe in Sub Pop 4 than in Sub Pop5 (Figure 3A-D). This indicated a close relationship between these two subsets of cells which might represent two states of the differentiation path of *Dullard* mutated cardiac NCC.

Strikingly, the genes that were differentially expressed between Sub Pop 3 and the Sub Pop 4/5 encode for known regulators of cell adhesion and epithelial-mesenchymal transition. On the one hand, in both Sub Pop 4 and Sub Pop 5 the levels of mesenchymal markers (*Snai2, Twist1, Cdh2, Mmp14, Rac1)* were lower than in the Sub Pop 3. In control embryos, the NCC expressing these markers were found nearby the endocardium, as demonstrated by an ISH for Snai2 or immunolabeling Twist1 (Figure 3E - Figure supplement 4B). In mutants, these pro-epithelial-mesenchymal transition factors were barely detected in the NCC (Figure 3E - Figure supplement 4B). This is providing a molecular mechanism for why NCC in Dullard mutants are very condensed and do not exhibit a mesenchymal shape nearby the endocardium (Figure 2E-G). On the other hand, cells in Sub Pop 4 and 5 displayed higher levels of the epithelial marker *Cdh5* and of *Sema3c*, the ligand of a signalling cascade regulating cohesive/metastatic balance in several cancers (Tamagnone, 2012), compared to SubPop3 cells. *In situ* hybridization of *Sema3c* further confirmed these results (Figure 3F - Figure supplement 4A,B). At distal OFT levels, this gene was expressed in scattered NCC in cushions of control embryos, while it was found in almost all NCC in mutants. Importantly, we observed a graded decrease of its expression along the OFT axis in both control and mutant embryos, which was reminiscent of the BMP response and condensation gradients described previously (Figure 3F).

In conclusion, our results show that Dullard regulates mostly cell autonomously the transcriptome of cardiac NCC, prevents the establishment of an epithelial-like state and promotes the expression of pro-mesenchymal genes.

## Discussion

By investigating the function of the Dullard phosphatase during cardiac NCC-mediated OFT septation, we have uncovered that the BMP-dependent condensation of the cardiac NCC is spatially regulated and sets the timing of cardiac cushions fusion. We showed that a gradient of BMP activation exists in the cardiac NCC along the OFT axis, and that its magnitude is under the control of the phosphatase *Dullard*. Yet, the establishment of this gradient is dependent on other mechanisms, as the gradient is still observed in absence of *Dullard* in the cardiac NCC. The establishment of this gradient is unlikely resulting from a corresponding gradient of ligand that would diffuse from a localized distal source, as described in other contexts (Bier & De Robertis, 2015). In fact, BMP4 is homogeneously expressed in the myocardium throughout the entire length of the OFT and not solely at the distal level (Jia et al., 2007; Jiao et al., 2003; Jones, Lyons, & Hogan, 1991; Liu et al., 2004; Zhang et al., 2010). Rather, intracellular signalling inhibitors could allow a temporal adaptation of cardiac NCC to BMP signals along with their migration towards proximal levels of the OFT. Prominent expression of *Smad6*, a BMP negative feedback effector, and of the diffusible inhibitor *Noggin*, are indeed observed in the OFT from E10.5 throughout great arteries formation and thus represent promising candidates (Choi, Stottmann, Yang, Meyers, & Klingensmith, 2007; Galvin et al., 2000).

Importantly, the BMP signalling gradient provides a means by which the zipper-like septation of the OFT can proceed. The levels of BMP stand as a sufficient parameter to tune the degree of condensation of cardiac NCC to the endocardium. In accordance, lowering BMP levels prevents the fusion of the cardiac cushions and leads to persistent truncus arteriosus (Jia et al., 2007; Stottmann et al., 2004), while increasing BMP downstream signalling triggers premature cardiac NCC condensation to the endocardium and accentuated OFT septation (Figure 2). This model is further supported by evidence showing that altering the ability of cardiac NCC to contract, migrate and adhere, hence to condensate, prevents the correct formation and positioning of the aorticopulmonary septum (Luo, High, Epstein, & Radice, 2006; Phillips et al., 2013; Plein et al., 2015).

Our study raises the question about the nature of the regulatory relationship existing between BMP signalling and *Sema3c* expression. Embryos from the *Pax3^Cre^* driver line harbour myogenic recombined cells which, as the cardiac NCC, show a striking increase in phosphorylation of Smad1/5/8 and *Sema3c* expression when *Dullard* is deleted (data not shown). This suggests that the regulatory influence of BMP signaling on *Sema3c* expression is not restricted to the context of cardiac NCC but also to other cell types. However, the Smads direct action on *Sema3c* expression remains unclear. Gata6, a member of the zinc finger family of transcription factors, has been described as an activator of *Sema3c* expression in the cardiac NCCs (Kodo et al., 2009; Lepore et al., 2006). Yet, we did not observe any significant difference of *Gata6* expression in *Dullard* deleted cardiac NCC (data not shown).

We have uncovered part of the cellular and molecular mechanisms by which BMP controls cardiac NCC behaviour. Upon Dullard deletion, over-activation of BMP signalling triggers a downregulation of mesenchymal markers reminiscent of a cardiac NCC transition towards epithelial-like states, with a loss of migratory freedom and increased cohesiveness between cells (Kim, Jackson, & Davidson, 2017). Furthermore, out of the transcriptional changes induced by the loss of *Dullard*, several lines of evidence support the idea that the up-regulation of *Sema3c* is likely to play a predominant role in the increased condensation of the cardiac NCC. In fact, the expression of *Sema3c* in the cardiac NCC is required for their convergence to the endocardium and was also shown to promote the aggregation of cardiac NCC in primary cultures as well as cancer cells in vivo (Delloye-Bourgeois et al., 2017; Feiner et al., 2001; Kodo et al., 2017; Plein et al., 2015; Toyofuku et al., 2008).

## Experimental procedures

### Mouse strains

All animal experiments were approved by the Animal Ethics Committee of Sorbonne University. We used the mouse strains described in the following papers with their MGI IDs: *Dullard^flox/flox^* ((Sakaguchi et al. 2013); in these mice exons 2 to 4 are floxed), *Pax3^Cre^* ((Engleka et al., 2005), MGI: 3573783), *Wnt1^Cre^* ((Danielian et al., 1998), MGI:2386570), *Rosa26mTmG* ((Muzumdar et al., 2007), MGI: 3716464), and *C57BL/6JRj* (Janvier Labs).

### Immunohistochemistry and imaging

Mouse embryos were collected at E11.5 and dissected in cold PBS, incubated 5min in 200mM KCl to stop heart beating and fixed for 2-3 hours in 4% PFA (Electron Microscopy Science, #15710S) at 4°C.

For immunostaining on cryosections, embryos were cryoprotected in 20% sucrose overnight at 4°C, embedded in OCT, and cryosectioned at 12µm thickness. Sections were permeabilized 10min in PBS/0.5% Triton, incubated for 1h in blocking buffer (5% goat serum in PBS) and overnight in primary antibody solution (in 1% BSA in PBS). After thorough washing in PBS they were incubated 1h in secondary antibody solution (in 1% BSA in PBS). Immunostainings were acquired using a Nikon Ti2 microscope, driven by Metamorph (Molecular Devices), equipped with a motorized stage and a Yokogawa CSU-W1 spinning disk head coupled with a Prime 95 sCMOS camera (Photometrics), then assembled and analyzed on Fiji (Schindelin et al., 2012). For wholemount staining, we followed the 3Disco protocol (Belle et al., 2017) to immunostain, clear and image with a lightsheet ultramicroscope (LaVision BioTec). Alternatively, micro-dissected hearts immunostained as described in (Belle et al., 2017), were clarified with BABB and imaged using a LSM700 confocal microscope (Carl Zeiss) (Gopalakrishnan et al., 2015). 3D renderings were generated using the Imaris software.

The primary antibodies used were raised against: GFP (chicken, Aves Labs, GFP-1020, 1/500), Pecam (rat monoclonal, Santa-Cruz Biotechnology, sc-18916, 1/200), Phospho-Smad1/5/8 (rabbit monoclonal, Cell Signalling Technology, 13820S, 1/500), myosin heavy chain, MyHC (mouse monoclonal, DSHB, MF20,1/300), Phospho-Histone H3 (rabbit, Cell Signalling Technology, 9701, 1/500), Cleaved Caspase-3 (rabbit monoclonal, Cell Signalling Technology, Asp175, 1/500). Secondary antibodies were bought from life technologies and were Donkey Igg coupled to Alexa fluorophores.

### In situ hybridisation

The *Sema3c* probe was provided by the lab of S. Zaffran (Bajolle et al., 2006). *In situ* hybridization on cryosections were processed following the protocol described in (Chotteau-Lelièvre, Dollé, & Gofflot, 2006).

### RNAscope

RNAscope® probes for *Dullard* (#456911), *Sema3c* (#441441), *Twist1* (#414701) and *Snai2* (#451191) were obtained from Advanced Cell Diagnostics, Inc. In situ hybridization was performed using the RNAscope® V2-fluorescent kit according to manufacturer’s instructions (Wang et al., 2012). For sample pre-treatments: H2O2 treatment was performed during 10 min at RT, retrieval 2 min at 98°C and slides were digested with Protease Plus reagent for 15 min at 40°C. After the probe detection steps immunostaining was performed as described above with fluorescent secondary antibodies. Sections were imaged using a 40x objective on a LSM700 microscope (Zeiss).

### Image analyses, quantification, statistical analysis

Mean levels of PSmads in the cardiac NCC were quantified using Image J thanks to a mask established on the GFP channel. Distances between cardiac NCC and between cardiac NCC and the endocardium, as well as the angle of the major axis of NCC to an axis linking the Ao and Pa poles of the endocardium were measured using Metamorph (Molecular Devices) and a home-made algorithm on Excel. Statistical analysis was performed with the Student’s t-test or Mann-Whitney test depending on normality. The analysis was performed using Prism Software (GraphPad). Statistical significance is represented as follows: ***p < 0.001. All results are shown as mean ± standard deviation.

### Cell culture and transfection

C2C12 cells were cultivated at 37°C/5% CO2, in growth medium (DMEM, 4.5g/L, D-glucose, 4mM L-glutamine, 1mM sodium pyruvate, 10% fetal calf serum). Plasmid transfection was performed using Lipofectamine 2000 (Life Technologies). 24 hours after transfection, BMP2 recombinant human protein (Thermo Fisher Scientific, #PHC7145) was applied for 1h on cells. Cells were fixed with 4% PFA and immunostained with the phospho-Smad1/5/8 antibody. Or, cells were washed in PBS, collected with PBS 1% SDS and passed through Qiashredder columns (Qiagen) to disrupt nucleic acids. Proteins extracts were then processed for Western blotting using pre-cast gels (Life Technologies) and transferred on nitrocellulose by semi-dry transfer (Bio-Rad).

### Tissue dissociation and FACS sorting

Mouse embryos were collected at E11.5 and placed in HBSS/1% FBS (HBSS +/+, Invitrogen) during genotyping. OFT were micro-dissected and dissociated by 15 min incubation in collagenase (0.1mg/ml in HBSS, C2139 Sigma) and thorough pipetting. HBSS (10% FBS) was added to the cells medium to stop the enzymatic reaction. OFT cell suspensions were centrifuged and resuspended in HBSS (1% FBS) before immunostaining. The panel of conjugated antibodies used for FACS included Ter119 Pe-Cy7 (Erythroid Cells, anti-mouse, Sony, Catalog No. 1181110, 1/300), CD45 APC-Cy7 (Rat anti-mouse, BD Pharmingen, Catalog No. 557659, 1/300), CD31 APC (Rat anti-mouse, BD Pharmingen, Clone MEC 13.3 Catalog No. 561814, 1/300) diluted in HBSS (1% FBS). Cells were centrifuged and resuspended in the antibody solution for a 25min incubation period (4°C, dark), washed 3 times, filtered (Fisher cell strainer, 70μm mesh) and 7AAD PE-Cy7 (1/800) was added in the cells suspension to exclude dead cells. Cells were sorted in a BD FACSAria™ III into 96-well plates loaded with RT-STA reaction mix (CellsDirect™ One-Step qRTPCR Kit, Invitrogen) and 0.2x specific TaqMan ® Assay mix (see Table 1 for assays list).

### Single-Cell gene expression

We proceeded as described in (Valente et al., 2019). Cells were sorted in RT-STA reaction mix from the CellsDirect™ One-Step qRT-PCR Kit (Life Technologies), reverse transcribed and specific target pre-amplified (20 cycles), according to the manufacturer’s procedures. Pre-amplified samples were diluted 5x with low EDTA TE buffer prior to qPCR analysis using 48.48 Dynamic Array™ IFCs and the BioMark TM HD System (Fluidigm). The same TaqMan® gene expression assays (20x, Life Technologies) were individually diluted 1:1 with 2x assay loading reagent (Fluidigm). Pre-amplified samples were combined with TaqMan Universal Master Mix (Life Technologies) and 20x GE sample loading reagent (Fluidigm). Loading of the 48.48 Dynamic Array TM IFCs and qPCR cycling conditions followed the Fluidigm procedure for TaqMan ® gene expression assays.

### Bioinformatic analysis

The analytic framework used followed the one described in (Perchet et al., 2018; Valente et al., 2019). It included notably a normalization of all the cycle thresholds (Ct) values extracted from the Biomark chips using the mean value of *Actb* and *Gapdh* housekeeping genes (Figure supplement 3A). For visualisation of the single cell multiplex qPCR, done on 44 genes, we generated a heatmap using the pHeatmap (v1.0.10) R package (v3.2.2). For unsupervised clustering, we used PhenoGraph that takes as input a matrix of N single-cell measurements and partitions them into subpopulations by clustering a graph that represents their phenotypic similarity. PhenoGraph builds this graph in two steps. First, it finds the k nearest neighbors for each cell (using Euclidean distance), resulting in N sets of k-neighborhoods. Second, it operates on these sets to build a weighted graph such that the weight between nodes scales with the number of neighbors they share. The Louvain community detection method is then used to find a partition of the graph that maximizes modularity. Given a dataset of N d-dimensional vectors, M distinct classes, and a vector providing the class labels for the first L samples, the PhenoGraph classifier assigns labels to the remaining N_L unlabeled vectors. First, a graph is constructed as described above. The classification problem then corresponds to the probability that a random walk originating at unlabeled node x will first reach a labeled node from each of the M classes. This defines an M-dimensional probability distribution for each node x that records its affinity for each class. PhenoGraph is implemented as Rphenograph (v0.99.1) R package (v3.2.3). We used boxplot for gene expressions of clusters obtained with PhenoGraph algorithm, from package ggplot2 (v3.1.0) R package (v3.2.2). For visualization and pathway analyses (tree), Destiny (v2.6.1) on R package (v3.6) generates the diffusion maps. Destiny calculates cell-to-cell transition probabilities based on a Gaussian kernel with a width σ to create a sparse transition probability matrix M. For σ, Destiny employs an estimation heuristic to derive this parameter. Destiny allows for the visualization of hundreds of thousands of cells by only using distances to the k nearest neighbors of each cell for the estimation of M. An eigen decomposition is performed on M after density normalization, considering only transition probabilities between different cells. The resulting data-structure contains the eigenvectors with decreasing eigenvalues as numbered diffusion components (DC), the input parameters and a reference to the data. These DC are pseudotimes identifying differentiation dynamics from our sc-qPCR data. Impact of gene expressions creating the different dimensions is represented as horizontal box plots, showing cells up- or down-regulating indicated genes in the plot.

## Acknowledgments

We thank the Cadot, Bitoun and Ribes Laboratories for discussions, Sigolène Meilhac for her help on understanding heart development concepts, Edgar Gomes laboratories, Isabelle Le Roux and Pascale Gilhardi-Hebenstreit for useful comments, Stéphane Zaffran for the *Sema3c* ISH probe and Morgane Belle for Lightsheet microscopy. The *Pax3^Cre^* and *Rosa26mTmG* transgenic lines were kindly provided by F. Relaix, and the *Wnt1^Cre^* line by A. Pierani. This work was supported by Agence Nationale pour la Recherche (ANR-14-CE09-0006-04) to BC; Association Institut de Myologie to BC. V.R. is an INSERM researcher, and work in her lab is supported by CNRS/INSERM ATIP-AVENIR program, as well as by a Ligue Nationale Contre le Cancer grant (PREAC2016.LCC). MV has a postdoctoral fellowship from the Laboratoire d’Excellence Revive (Investissement d’Avenir; ANR-10-LABX-73).

## Authors contributions

JFD, PM, GC, MV and BC performed the experiments; MP analyzed the transcriptomics data; JFD, VR and BC conceived and designed the experiments; JFD, VR and BC wrote the manuscript with assistance from MV and GC; DSO and RN provided reagents.

---

**Figure supplement 1:**
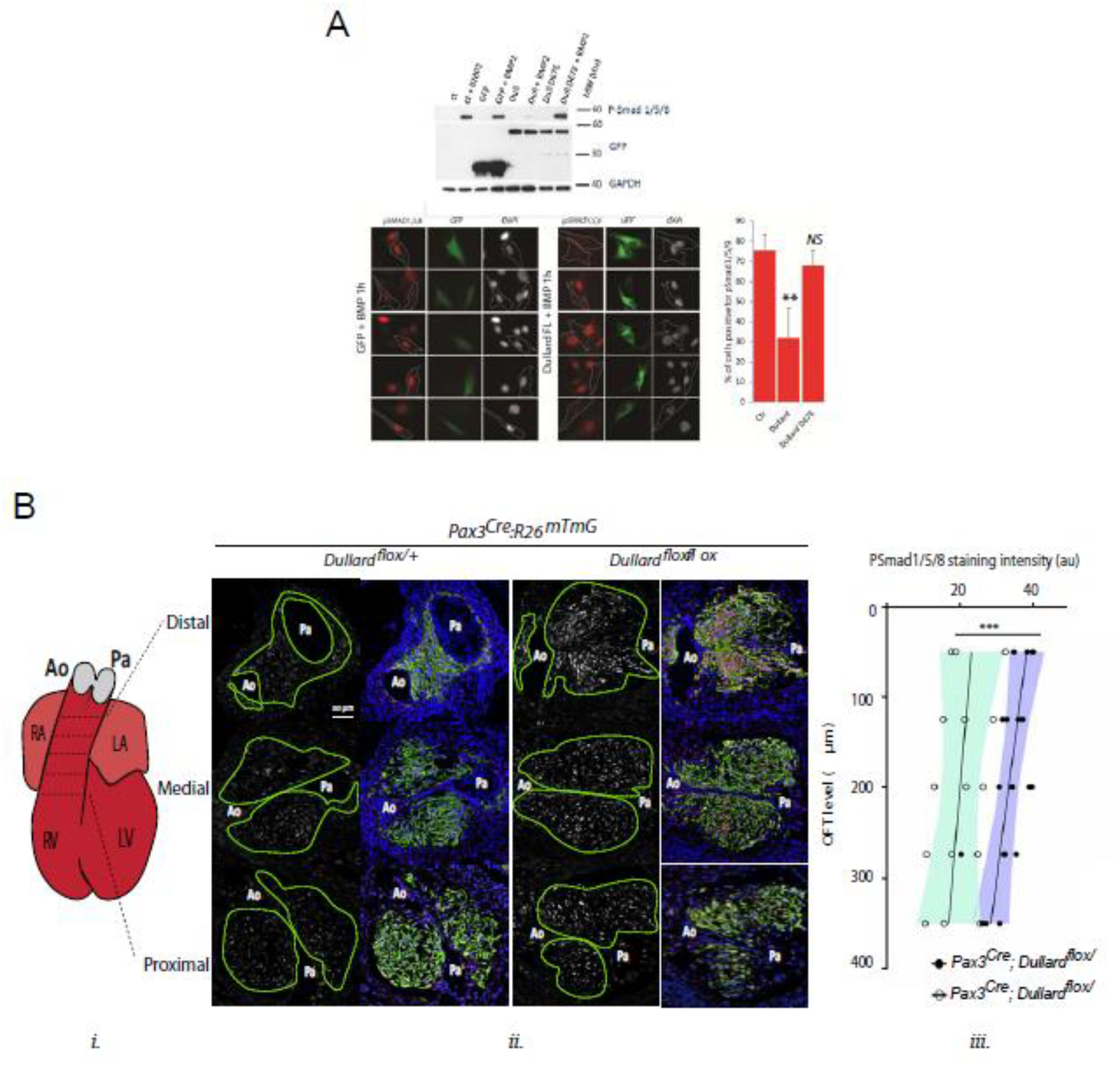
Dullard phosphatase is a negative regulator of BMP signalling in mammalian cells. **(A)** Top: Western blot detecting the phosphorylated forms of Smad1/5/8, GFP or Gapdh in C2C12 muscle cells exposed with or without BMP2 for 1h. These cells were non transfected (ct) or transfected with either a GFP expressing plasmid (GFP), a GFP tagged version of the wild-type Dullard (Dull) or of Dullard carrying D67E mutation in its phosphatase domain (Dull D67E). Dullard inhibits BMP2 mediated phosphorylation of Smad1/5/8; this inhibition is dependent on the functionality of its phosphatase domain. Bottom-left: Immunofluorescence for PSmads and GFP and DAPI staining in C2C12 transfected with GFP or GFP-Dullard and exposed for 1 h to BMP2 showing that only cells transfected with Dullard do not have nuclear phosphorylated Smad1/5/8. Right: Quantification of the number of PSmads positive C2C12 cells exposed to BMP2 for 1h and transfected with GFP (ct), Dullard or Dullard carrying the D67E mutation (n=3 independent experiments; Student t-test **: p-value < 0.01. **(B) Bi.** Schematics of E11.5 heart showing the position of the transverse sections used to quantify the levels of the phosphorylated forms of Smad1/5/8 (PSmads) in iii. **Bii.** Immuno-labelling for PSmads and GFP and DAPI staining on transverse sections across the OFT at 3 distinct distal-proximal levels in E11.5 of control Pax3^Cre^; Dullard^flox/+^; Rosa26mTmG and mutant Pax3^Cre^; Dullard^flox/flox^; Rosa26mTmG hearts. Pale green dotted lines delineate the area colonized by cardiac NCC. **Biii.** Quantification of PSmads levels in cardiac NCC along the distal-proximal axis of the OFT of E11.5 embryos with the indicated genotype (dots: values obtained on a given section; n>4 embryos per genotype recovered from at least 3 liters; the black line is the linear regression, the coloured areas delineate the 95% confidence intervals, ***: p-value < 0,001 for a two way-Anova statistical test). Ao: aortic artery, Pa: pulmonary artery.

**Figure supplement 2:**
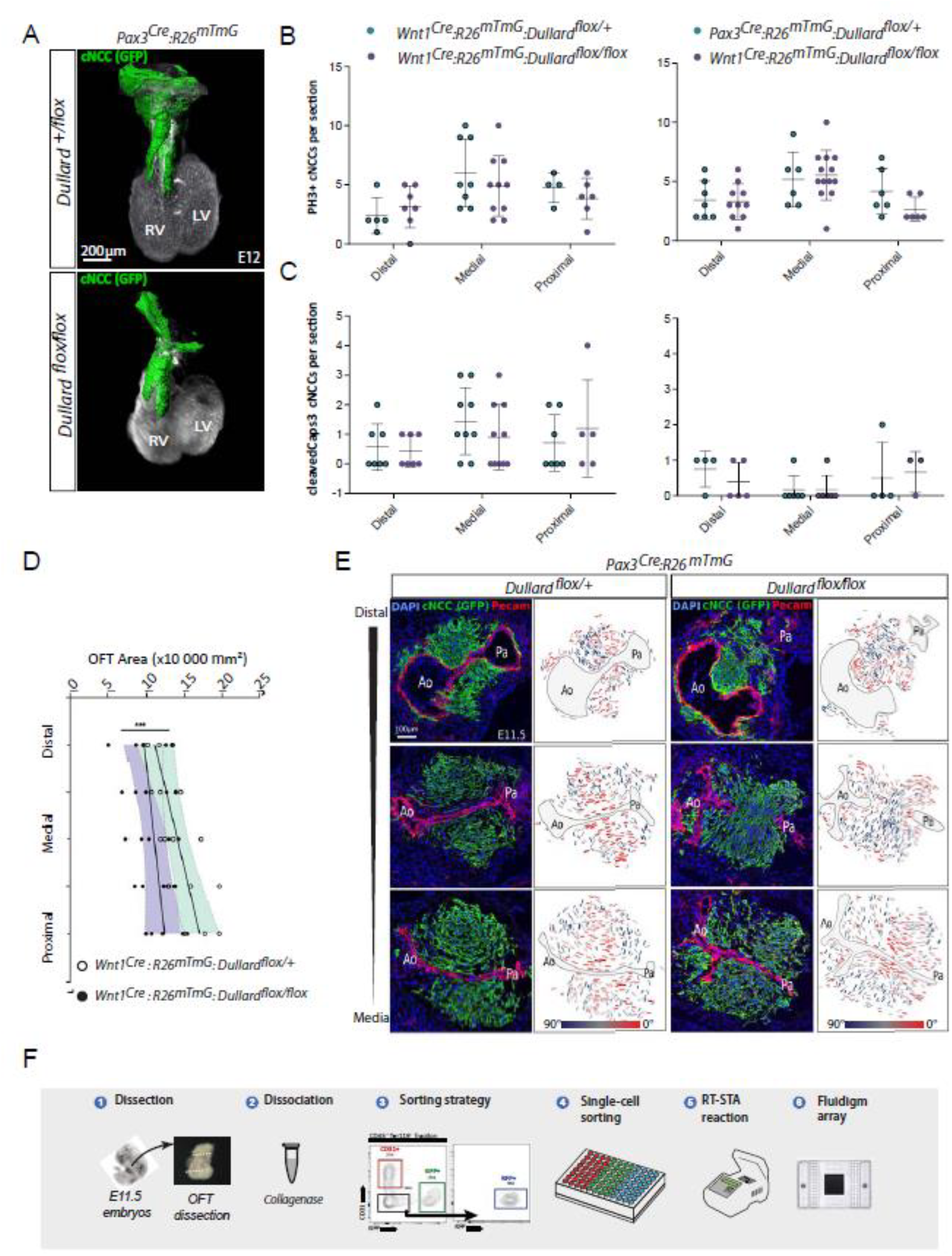
Morphogenetic defects in Dullard NCC mutants. **(A)** Three-dimensional rendering of cardiac NCC (green) over Pecam (white) after BABB clearing and confocal acquisition of whole E11.5 hearts isolated from embryos with the indicated genotype (n=2 per genotype). **(B, C)** Quantification of PH3^+^ mitotic (B) and activated caspase 3^+^ apoptotic cardiac NCC per OFT section at the indicated distal-proximal OFT axis levels (dots: value per section; bars: mean±s.d.; n.s.: non-significant differences evaluated using a Mann-Whitney test). **(D)** Quantification of the OFT surface area measured at distinct distal-proximal axis levels in control and Dullard mutants at E11.5 (dots: values obtained on a given section; n>4 embryos per genotype recovered from at least 3 liters; the black line is the linear regression, the coloured areas delineate the 95% confidence intervals, ***: p-value < 0,001 for a two way-Anova statistical test). **(E) Ei-vi.** Immuno-labelling for Pecam (red) and GFP (green) and DAPI (blue) staining on transverse sections along the distal-proximal axis of the OFT in E11.5 embryos with the indicated genotypes (n>10 embryos collected from more than 3 liters). Brackets in i and iv highlight the symmetric and asymmetric Ao and Pa poles in control and mutant embryos, respectively. Arrowheads in ii and v point at the unruptured and ruptured in control and mutant embryos, respectively. **Ei’-vi’.** Orientation of the major axis of NCC cells relative to Ao-Pa axis colour coded as indicated in the section shown in Di-vi. **(F)** Experimental steps performed to profile gene expression in OFT single-cells sorted from five Wnt1^Cre^; Dullard^flox/+^; Rosa26mTmG and five Wnt1^Cre^; Dullard^flox/flox^; Rosa26mTmG E11.5 embryos. At least 70 cells were isolated per gate and genotype (GFP+, CD31+, RFP+).

**Figure supplement 3:**
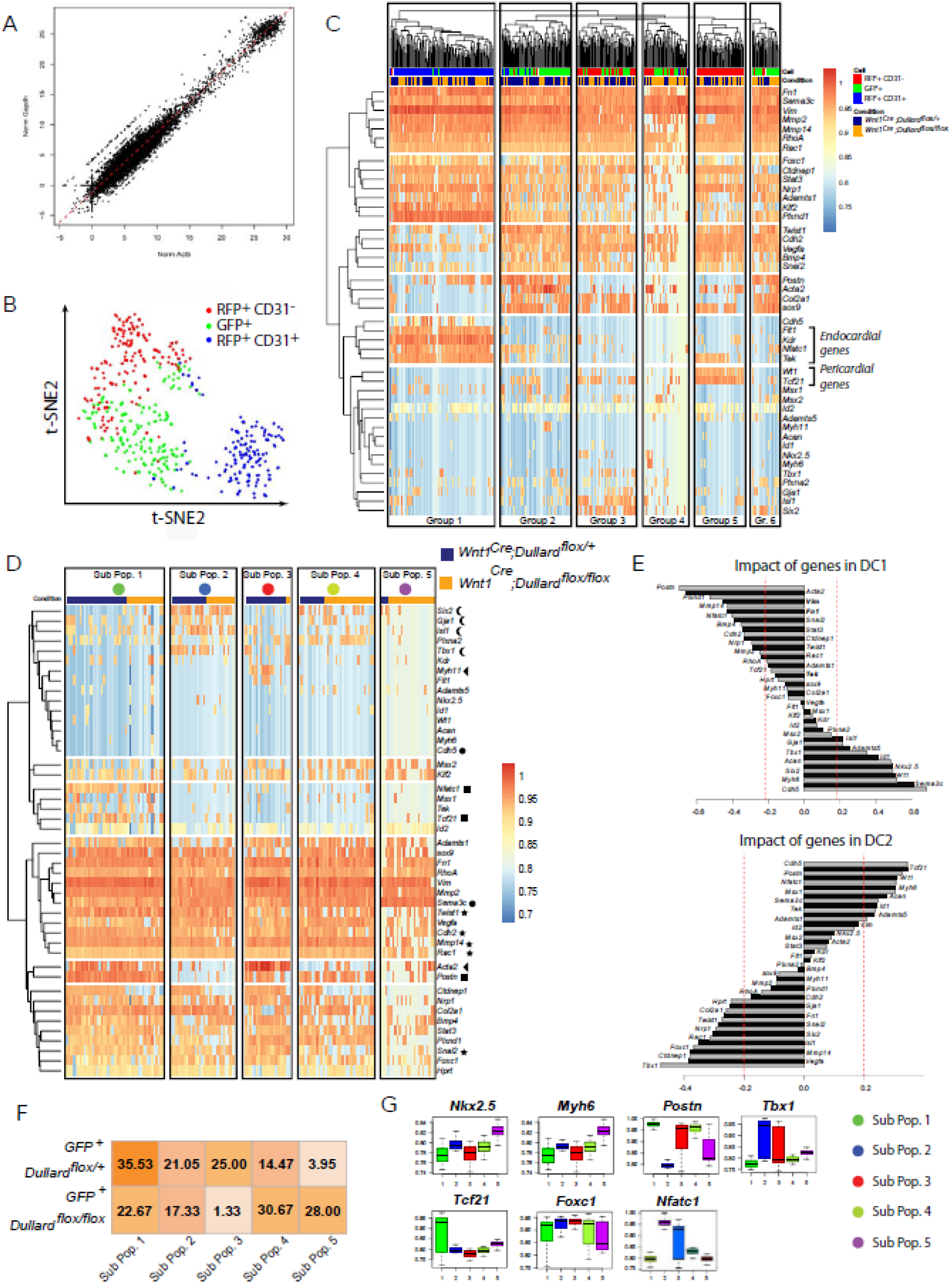
Single-cell transcriptional analyses of OFT cells at E11.5. **(A)** Graph showing the distribution of all cells analyzed (dots) as a function of normalized expression values of the house keeping genes Actb and Gapdh and a linear regression (black line). **(B)** tSNE plot showing the distribution of 44-genes-based transcriptomes of 433 OFT cells expressing the indicated markers isolated from both Wnt1^Cre^; Dullard^+/flox^; Rosa26mTmG and Wnt1^Cre^; Dullard^flox/flox^; Rosa26mTmG E11.5 embryos. **(C)** Unsupervised clustering heatmap of the 433 OFT isolated cells from Wnt1^Cre^; Dullard^+/flox^; Rosa26mTmG and Wnt1^Cre^; Dullard^flox/flox^; Rosa26mTmG based on the gene expression level of the 44 genes included in the panel (Table S1). Six different groups of cells can be discriminated, among which the endocardial (Group1) cells expressing high levels of Flt1, Kdr, Nfatc1 and Tek, and the epicardial (Group5) cells expressing high levels of Wt1, Tcf21. **(D)** Unsupervised clustering heatmap of 151 GFP+ NCC isolated from Wnt1^Cre^; Dullard^+/flox^; Rosa26mTmG and Wnt1^Cre^; Dullard^flox/flox^; Rosa26mTmG based on the gene expression level of the 44 genes included in the panel (Table S1). Five different sub-populations can be identified, two of which (Sub-Pop1 and 2) are not Dullard dependent, while the three others (Sub-Pop3 to 5) are enriched for either mutant (Sub-Pop4,5) or control cells (Sub-Pop3). Stars indicates genes involved in the mesenchymal state. The circles marks instead the genes associated with an epithelial fate. The arrows points at genes encoding for smooth muscle specific genes marking the Pop3. Crescent point to typical cardiac genes, whereas squares point to genes involved in valves formation. **(E)** Genetic composition of the diffusion components (DC) 1 and 2, on which are based the diffusion maps presented in Figure supplement 3E. **(F)** Fraction of mutant and control cells present in all 5 sub-populations defined in the unsupervised clustering in Figure supplement 3E. **(G)** Boxplot representation of the expression levels of genes highlighted in DC1 and 2 (Figure supplement 3D) and not presented in Figure 3D, in the GFP+ NCC cells from both genotypes (mean±s.d.).

**Figure supplement 4:**
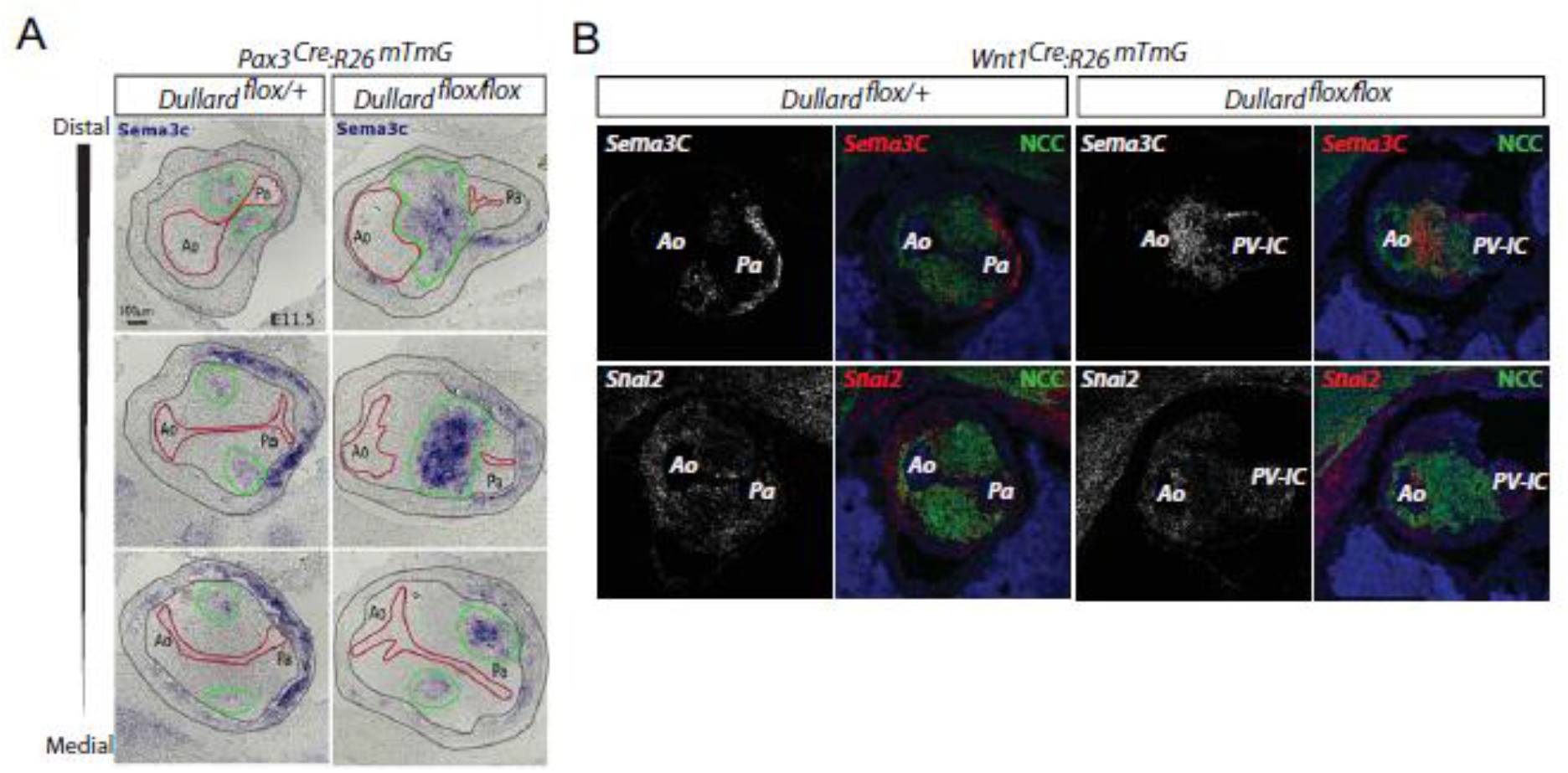
Dullard loss in cardiac NCC alters the expression of key epithelial and mesenchymal traits drivers. **(A)** *Sema3c* expression assessed by ISH on transverse sections from distal to medial OFT levels of E11.5 *Pax3^Cre^; Dullard^flox/+^* and *Pax3^Cre^; Dullard^flox/flox^* embryos. The endocardium is delineated with a red line, the cardiac NCC areas with a green line and the myocardium with grey lines. **(B)** *Sema3c* and *Snai2* mRNA distribution assessed by RNAsecope, GFP immuno-labelling and DAPI staining in transverse sections of E11.5 control *Wnt1^Cre^; Dullard^flox/+^*and mutant *Wnt1^Cre^; Dullard^flox/flox^* OFTs at distal (**Bi**) and medial levels (**Bii**). Ao: Aorta; Pa: pulmonary artery

**Table supplement 1:**
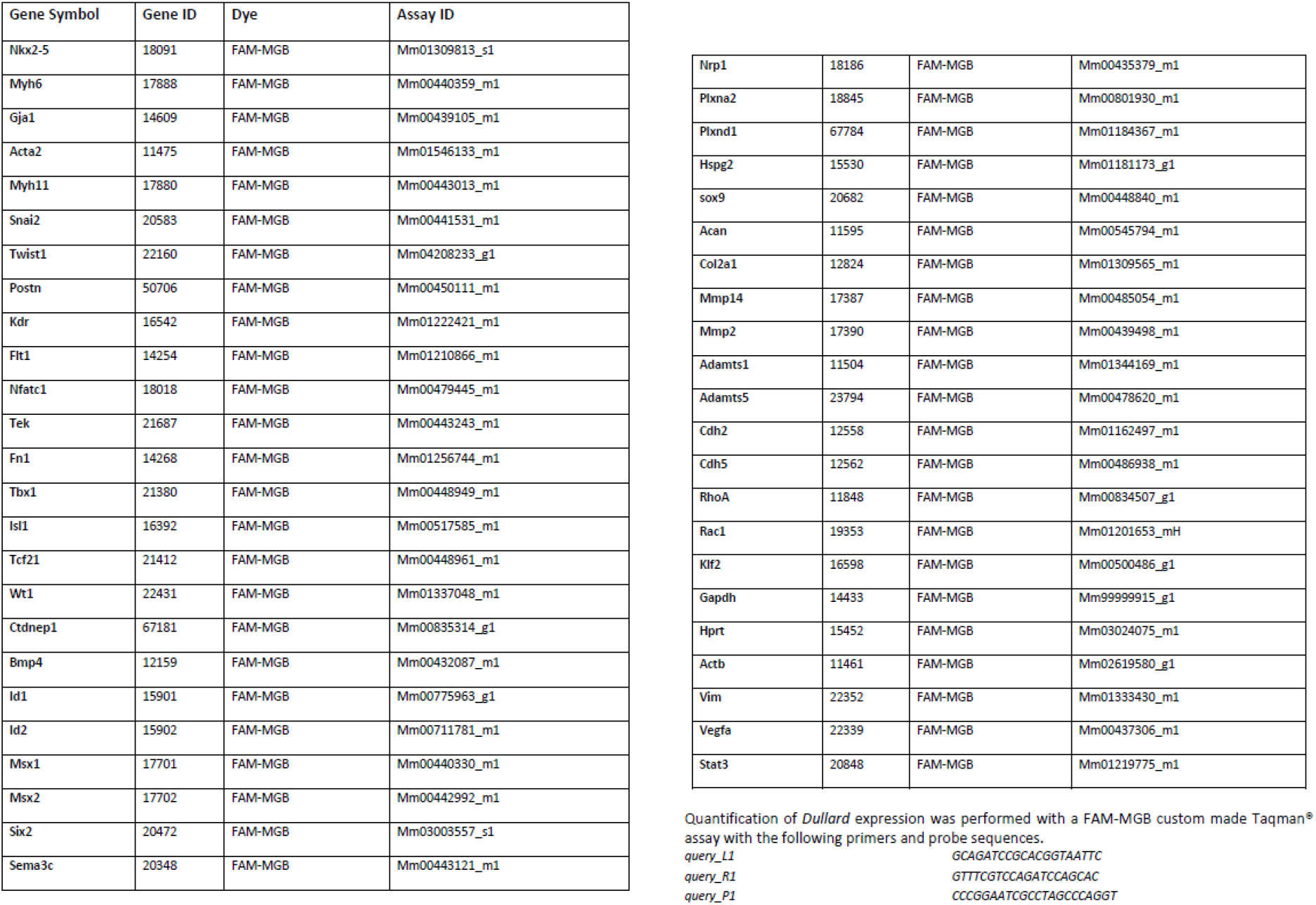
List of TaqMan® gene expression assays (20x, Life Technologies) used for single-cell qPCRs experiments.

**Movie supplement 1/2:** Three-dimensional rendering of cardiac NCC (green isosurface) over Pecam (red isosurface) after 3Disco clearing and Lightsheet acquisition of *Pax3^Cre^*; *Dullard^flox/+^; Rosa26mTmG*and *Pax3^Cre^*; *Dullard^flox/flox^; Rosa26mTmG*E12 embryos. An atrophy of the pulmonary artery is observed.

**Movie supplement 3/4:** Three-dimensional rendering of cardiac NCC (green isosurface) over Pecam (white) after 3Disco clearing and Lightsheet acquisition of *Pax3^Cre^*; *Dullard^flox/+^; Rosa26mTmG* and *Pax3^Cre^*; *Dullard^flox/flox^; Rosa26mTmG* E12 embryos. No defect in the OFT colonization of mutant cardiac NCC is observed.

**Movie supplement 5/6:** Three-dimensional rendering of cardiac NCC (green) over Pecam (white) after BABB clearing and confocal acquisition of *Wnt1^Cre^*; *Dullard^flox/+^; Rosa26mTmG* and *Wnt1^Cre^*; *Dullard^flox/flox^; Rosa26mTmG* E11.5 embryos. No defect in the OFT colonization of mutant cardiac NCC is observed.

